# Development of cross-orientation suppression and size tuning and the role of experience

**DOI:** 10.1101/201228

**Authors:** Marjena Popović, Andrea K. Stacy, Mihwa Kang, Roshan Nanu, Charlotte E. Oettgen, Derek L. Wise, József Fiser, Stephen D. Van Hooser

## Abstract

Many sensory neural circuits exhibit response normalization, which occurs when the response of a neuron to a combination of multiple stimuli is less than the sum of the responses to the individual stimuli presented alone. In the visual cortex, normalization takes the forms of cross-orientation suppression and surround suppression. At the onset of visual experience, visual circuits are partially developed and exhibit some mature features such as orientation selectivity, but it is unknown whether cross-orientation suppression or surround suppression are present at the onset of visual experience or require visual experience for their emergence. We characterized the development of these properties and their dependence on visual experience in ferrets. Visual experience was varied across three conditions: typical rearing, dark rearing, and dark rearing with daily exposure to simple sinusoidal gratings (14-16 hours total). Cross-orientation suppression and surround suppression were noted in the earliest observations, and did not vary considerably with experience. We also observed evidence of continued maturation of receptive field properties in the second month of visual experience: substantial length summation was observed only in the oldest animals (postnatal day 90); evoked firing rates were greatly increased in older animals; and direction selectivity required experience, but declined slightly in older animals. These results constrain the space of possible circuit implementations of these features.

**Significance Statement:** The development of the brain depends on both nature – factors that are independent of the experience of an individual animal – and nurture – factors that depend on experience. While orientation selectivity, one of the major response properties of neurons in visual cortex, is already present at the onset of visual experience, it is unknown if response properties that depend on interactions among multiple stimuli develop without experience. We find that the properties of cross-orientation suppression and surround suppression are present at eye opening, and do not depend on visual experience. Our results are consistent with the idea that a majority of the basic properties of sensory neurons in primary visual cortex are derived independent of the experience of an individual animal.

## Introduction

One of the most ubiquitous features of sensory receptive fields across species, modalities, and cortical hierarchies is the property of divisive normalization: cells exhibit responses to combinations of stimuli that are less than the sum of the responses to the individual stimuli (Heeger, 1992; Carandini et al., 1997; Tolhurst and Heeger, 1997; Simoncelli and Heeger, 1998; Britten and Heuer, 1999; Reynolds and Heeger, 2009; Olsen et al., 2010; Ohshiro et al., 2011; Ruff et al., 2016). In the primary visual cortex, the most-studied form of normalization is cross-orientation suppression, which occurs when the response of a neuron to an optimally oriented grating stimulus is suppressed by a superimposed orthogonal grating (plaid stimulus) that does not, by itself, elicit a response (Adelson and Movshon, 1982; Morrone et al., 1982; Morrone et al., 1987; DeAngelis et al., 1992). In addition, some models also posit that other contextual interactions – such as size tuning – are simply forms of normalization (Rubin et al., 2015). Size tuning includes surround suppression (Hubel and Wiesel, 1965; Gilbert, 1977; Bolz and Gilbert, 1986; DeAngelis et al., 1994) and length summation (Bolz and Gilbert, 1989; Chisum et al., 2003; Van Hooser et al., 2006). Despite the importance of normalization in sensory computation, it remains unknown whether the development of normalization manifested as cross-orientation suppression or size tuning requires sensory experience or, rather, is formed without sensory experience.

The proper development of most neural circuits is contingent on both experience-independent and experience-dependent factors. Some response properties of sensory neurons are present before sensory experience. At the onset of visual experience, neurons in carnivore V1 already exhibit tuning for stimulus orientation, spatial frequency, and temporal frequency, and this tuning is elaborated or altered by experience-dependent processes (Chapman and Stryker, 1993; White et al., 2001; Li et al., 2006). Other response properties require experience for their expression. For example, direction selectivity is very weak at the onset of visual experience, and develops rapidly when the ferret experiences moving visual stimuli (Li et al., 2006; Li et al., 2008; Van Hooser et al., 2012; Smith et al., 2015).

To study the influence of experience on cross-orientation suppression and size tuning, we compared visual receptive field properties in dark-reared animals and typically-reared animals at several ages. Additionally, we exposed some dark-reared animals to several hours of viewing artificial stimuli – large gratings of single orientations. These impoverished stimuli lacked the simultaneous presentation of multiple orientations, spatial, and temporal frequencies as well as variation in size typical of natural images. This allowed us to further explore the influence of the quality of visual experience on the emergence of cross-orientation suppression and size tuning.

We found that cross-orientation suppression and surround suppression were present in both dark-reared animals and in experienced animals, although the magnitude of this tuning varied slightly with experience and age. Cross-orientation suppression and surround suppression in animals whose experience was limited to large gratings of single orientations did not differ from that of typically reared animals, suggesting that experience is not critical for the emergence of these properties.

Finally, we uncovered unexpected evidence that basic visual cortical response properties continue to mature even after a month of visual experience. In typically-reared ferrets, direction selectivity reached a peak about a week after eye opening and was reduced later, suggesting that some receptive field features may change with age in a non-monotonic fashion. In addition, evoked firing rates and length summation increased substantially during the second month of visual experience. These changes occurred after the outgrowth of the long-range horizontal connections within visual cortex (Durack and Katz, 1996; Ruthazer and Stryker, 1996; White et al., 2001), but closely follow the growth of synaptic density to its peak at about 90 days of age (Erisir and Harris, 2003; White and Fitzpatrick, 2007).

## Materials and Methods

All experimental procedures were approved by the Brandeis University Animal Care and Use Committee and performed in compliance with National Institutes of Health guidelines.

### Experimental groups

Female sable ferrets (*Mustela putorius furo*) used in the experiment were split into five study groups (**Figure 1**):

**1 Dark-reared (n=5)**: The animals were reared with 3-5 littermates and their jill in complete darkness starting 1-3 days before eye opening (postnatal day 27-30) until the experiment at age P40-P42. The kits were nursed by their jill until spontaneous weaning; water and soft diet were available ad libitum. These animals were also used in an unrelated study; we took advantage of the opportunity to make additional discoveries with animals that were being initially studied for other purposes. As a part of this unrelated experiment, at P30 (with eyes still closed) the ferrets were implanted with chronic microelectrode arrays in left V1, and at P33 and P37 they were subject to two 80 min long, head-fixed recording sessions. Stimuli consisted of 20 minutes of a greyscale natural movie, 20 minutes of drifting gratings, 20 minutes of block noise and 20 minutes of a dark screen (Berkes et al., 2011). While it would have been ideal (for the study discussed here) to have had access to animals that did not include this procedure, our previous research has found that <3 hours of visual experience does not cause a substantial increase in direction selectivity, so we did not expect substantial influence of the experience that occurs during these measurements(Clemens et al., 2012; Roy et al., 2016; Ritter et al., 2017). During the post-operative recovery they were additionally fed milk replacement (KMR, PetAg) with a syringe. The need for additional syringe feeding was not related to dark rearing – it did not differ from that observed in typically-reared ferrets of corresponding age after surgery. Ferrets were observed and their weight was monitored daily using night vision goggles with an infrared light source. No effects of dark rearing on the animals’ general health and behavior were observed.

**2 Dark-reared, trained (n=7)**: The animals were reared with 3-5 littermates and their jill in complete darkness starting 1-3 days before eye opening (postnatal day 27-30) until the experiment at age P40-P42. As a part of the previously described unrelated experiment, at P30 they were implanted with chronic microelectrode arrays in left V1 and had headposts affixed, and at P33 and P37 they were subject to two 80 min-long recording sessions. Between P33 and P37 the ferrets were exposed to controlled visual experience. Awake ferrets were head-fixed and placed in front of a screen inside a dark box for two 1.5hr long sessions daily with a 1.5hr break in between the sessions adding to a total of 14-16 hours. The training stimuli consisted of bidirectionally drifting sinusoidal gratings of varying orientation (from horizontal to vertical in 22.5° steps) at 0.1 cycle/degree spatial frequency, 4Hz temporal frequency, and 100% contrast in 20 minute blocks separated with 10 min of mean luminance. Other than the daily training sessions the rearing conditions for ferrets in this experimental group were identical to those in the dark-reared group. Efforts were made to keep the dark-reared ferret kits awake during the period of visual stimulation (gentle tapping or gentle hand clapping), although these very young animals frequently dozed for several minutes during the visual exposure.

**3 Typically-reared, P40 (n=8)**: The animals were reared with 3-5 littermates and jill in a 12 h light/dark cycle environment until the experiment at age P40-P42. As a part of the previously described unrelated experiment, at P30 they were implanted with chronic microelectrode arrays in left V1 and had headposts affixed, and at P33 and P37 they were subject to two 80 min-long recording sessions.

**4 Typically-reared, P60 (n=4)**: The animals were reared in a 12 h light/dark cycle environment until the experiment at age P59-P66.

**5 Typically-reared, P90 (n=6)**: The animals were reared in a 12 h light/dark cycle environment until the experiment at age P90-P92.

**Figure 1.**
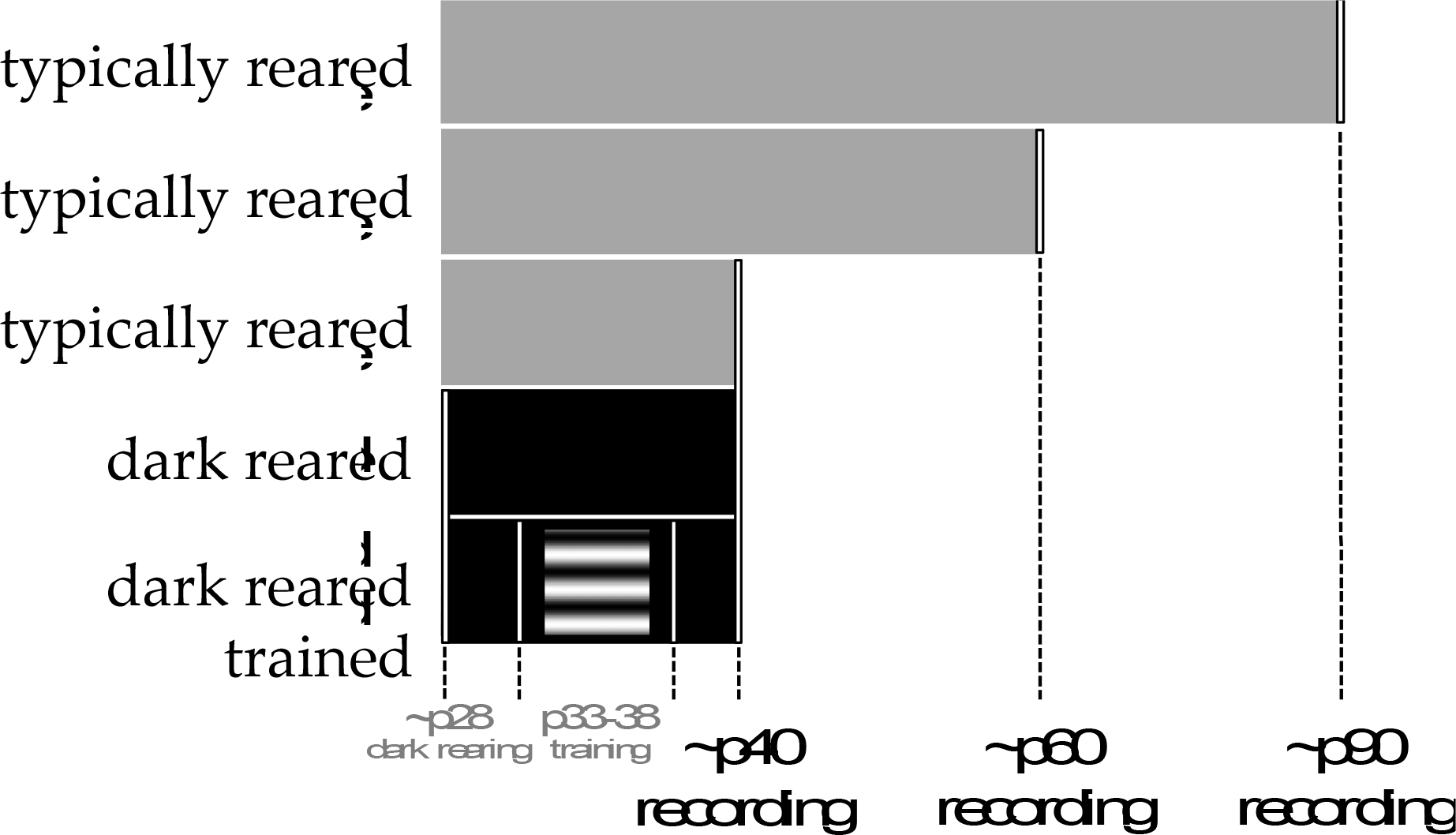
Experimental groups. We studied the development of receptive field properties while varying visual experience and age. We reared three groups of animals until approximately postnatal day 40 (P40, see Methods) according to different protocols. One group (typically-reared, P40) was provided 12 hours of normal light each day. Another group (dark-reared P40) was raised in darkness from P28 (about 3-5 days before normal eye opening) and had very impoverished visual experience. A third group (dark-reared and trained, P40) was raised in the dark from P28 but was provided with 14-16 total hours of experience with simple sinusoidal grating stimuli over several sessions. These groups were compared to animals with approximately 1 month of typical visual experience (typically-reared, P60) and approximately 2 months of visual experience (typically-reared P90).

### Survival surgery

All the ferrets in the dark-reared, dark-reared trained, and typically-reared ~P40 group were also used in an unrelated chronic recording experiment. For this experiment, ferrets had a 2×8 microwire electrode array implanted into V1 in the left hemisphere at age P30, while their eyes were still closed. Immediately before the surgery and up to 48 hours after surgery, ferrets were intramuscularly (IM) administered analgesic ketoprofen (1mg/kg) and antibiotic penicillin (27mg/kg) and orally administered analgesic tramadol (2mg/kg). During the surgery, ferrets were anesthetized with an intramuscular injection of ketamine-xylazine cocktail (20mg/kg and 2mg/kg, respectively), and given atropine (0.04mg/kg) to reduce secretions. Surgical margins were infused with 0.2 ml of the local analgesic bupivacaine. At the end of surgery, anesthesia was reversed using the xylazine antidote yohimbine (0.5mg/kg). Importantly, the last dose of analgesics was given 24hrs before the start of training. Body temperature was maintained at 37°C using a thermostatically controlled heating pad, heart rate was continuously monitored, and hydration was maintained throughout by subcutaneous injections of lactated Ringer’s solution (3 ml/kg/h). The cranium was exposed and a 4mm×4mm craniotomy made over V1. A durotomy was performed with a 31-gauge needle before placing the electrode array into the brain. The brain was sealed with a low toxicity silicone elastomere (Kwik-Cast, World Precision Instruments) and the electrode and headpost were affixed to the skull using 6 skull screws and light cured dental composite (Flow-It ALC, Pentron). After the animals were ambulatory, they were transferred back to the cage with their littermates and jill.

### Acute surgical procedures

Ferrets were sedated with ketamine (20 mg/kg IM). Atropine (0.16–0.8 mg/kg IM) was administered to prevent bradycardia and reduce bronchial and salivary secretion and dexamethasone (0.5 mg/kg IM) administered to reduce inflammation and swelling. The animal was deeply anesthetized with a mixture of isoflurane, oxygen, and nitrous oxide through a face mask while tracheotomy was performed. Once the tracheotomy was done, the animal was ventilated with 1–2% isoflurane in a 2:1 mixture of nitrous oxide and oxygen. A cannula was inserted into the intraperitoneal (IP) cavity for delivery of 5% dextrose in lactated Ringers solution (3 ml/kg/h). Body temperature was maintained at 37°C using a thermostatically controlled heating pad. End-tidal CO_2_ levels and respiration rate were monitored and kept within the appropriate physiological range (3.5-4%). The animal was held in place by a custom stereotaxic frame. All wound margins were infused with bupivacaine. Silicone oil was placed on the eyes to prevent damage to the cornea. A 4×4 mm craniotomy was made over V1 in the right hemisphere, and the dura was removed with a 31-gauge needle.

Before the start of recording, ferrets were paralyzed using a neuromuscular blocker (gallamine triethiodide 10 - 30 mg/h/kg), delivered through the IP cannula, in order to suppress spontaneous eye movements. The nitrous oxide to oxygen ratio was adjusted to 1:1. Adequate anesthesia was maintained by continuously monitoring the animals’ EKG and adjusting the isoflurane concentration. At the conclusion of the experiment the animal was killed and transcardially perfused to retrieve the brain for histology.

### Electrophysiological recordings

Carbon fiber electrodes (Carbostar-1, Kation Scientific) were used for all recordings. The signal was amplified using the RHD2000 amplifying/digitizing chip and USB interface board (Intan Technologies) and acquired and clustered using a Micro1401 acquisition board and Spike2 software (Cambridge Electronic Design, LLC). Spike sorting was done manually using Spike 2 software.

An electrode was inserted into the brain using a Sutter Instruments MP-285 manipulator. In order to reduce sampling bias, we recorded from any site that had a signal to noise ratio sufficient for isolation and had a response that appeared to be modulated by presentation of drifting gratings. Data are reported from all units that are responsive enough to be included in analysis (see below). After finishing the recording at one site, the electrode was lowered at least 40μm before attempting to identify a suitable subsequent recording site. The experiment was concluded once white matter was reached or once the animal’s physiological indicators became unstable.

### Visual stimulation

Visual stimuli were created in Matlab using the Psychophysics Toolbox (Brainard, 1997; Pelli, 1997) and displayed on a 21” flat face CRT monitor (Sony GDM-520) with a resolution of 800x600 and a refresh rate of 100Hz. The monitor was positioned 20cm away from the animals’ eyes, such that it was subtending 63°×63° of visual angle. For each unit we isolated, we first determined the ocular dominance and occluded the non-dominant eye. We then used circular patches of drifting sinusoidal gratings of varying sizes to manually map receptive fields. We moved the monitor to accommodate all eccentricities without varying the distance of the monitor from the animal.

### Immunohistochemistry

Upon completion of each experiment, electrolytic lesions were made at the final recording site and at ~300μm from the surface of the cortex to enable the reconstruction of the electrode track. Following a transcardial perfusion, the brain was placed in 4% paraformaldehyde in 0.1 M PBS at 4°C for 24 h, and then moved to 10% sucrose in PBS at 4°C for 24 – 48 h, followed by 30% sucrose in PBS at 4°C for 24 – 48 h.

### Data analysis

We recorded from a total of 335 V1 neurons of 30 female sable ferrets in 5 experimental conditions. Cells with a response rate below 2 spikes/sec were excluded during analysis, but were still recorded from if they appeared to be modulated by the stimuli during the experiment. The actual number of cells included in analysis varied across conditions (**Table 1**). Additional exclusion criteria for specific analyses are discussed in separate sections below.

**Table 1.** Number of cells analyzed per condition

|  | dark-reared<br>P40 | dark-reared<br>trained P40 | typically-reared P40 | typically-reared P60 | typically-reared P90 |
| --- | --- | --- | --- | --- | --- |
| ages | 2×40, 3×42 | 1×39, 2×40,<br>4×42 | 2×40, 2×41,<br>4×42 | 2×59, 1×61,<br>1×66 | 4×90, 1×91,<br>1×92 |
| # animals | 5 | 7 | 8 | 4 | 6 |
| # cells | 71 | 77 | 79 | 45 | 63 |
| # cells included |  |  |  |  |  |
| DS | 50 | 54 | 47 | 29 | 49 |
| SF | 66 | 73 | 53 | 43 | 59 |
| TF | 69 | 71 | 55 | 42 | 55 |
| contrast | 67 | 70 | 40 | 39 | 58 |
| size | 44 | 37 | 31 | 32 | 45 |
| Cross-orientation<br>suppression | 54 | 52 | 55 | 40 | 60 |

### Orientation and direction tuning

We characterized the orientation and direction selectivity of all cells using bidirectionally drifting sinusoidal grating stimuli of varying direction (22.5° steps) at 0.1 cycle/degree spatial frequency, 4Hz temporal frequency and 100% contrast. Orientation/direction tuning curves were fit with a mixture of two Gaussians in circular space, forced to peak 180° apart and to have the same width *σ*(Carandini & Ferster, 2000):

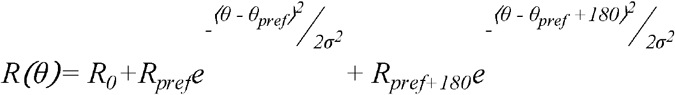

where *θ* is the stimulus direction in circular space (0°-360°), *R*_*0*_ is a constant offset, *θ*_*pref*_ is the preferred orientation, the tuning width, (*R*_*pref*_) is the increment above offset to the preferred direction, (*R*_*pref*+180_) is the increment above offset to the opposite direction, and the tuning width (half-width at half height) is given by 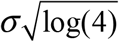.

To ensure good fitting, we constrained the fitting parameters: *σ ≥α/2*, where *α* is the stimulus angle step size (22.5°); −*R*_*max*_*≤R*_*0*_*≤R*_*max*_, where is *R*_*max*_ the highest observed response for any stimulus; *0 ≥R*_*prff*_, *R*_*pref+180*_ *≤ 3R*_*max*_. We initiated iterative fitting at parameter values expected to produce a good fit: 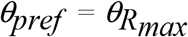; *R*_*pref*_ = *R*_*pref+180*_ = *R*_*max*_; *R*_*0*_ = *0*. We performed fitting for *σ ≡ (α/2, α, 40°, 60°, 90°)* and selected the fit with the lowest least squared error. Finally, we eliminated from further analysis cells that did not exhibit significant orientation tuning as quantified by Hotelling’s T^2^ test performed on orientation vector for each trial. This was done because the fitting method has been shown to produce large errors in *θ*_*pref*_ at low OI values (Mazurek et al., 2014).

Orientation selectivity was quantified using circular variance (Batschelet, 1981). Direction selectivity was quantified using direction index (DI), a normalized difference between the responses for the preferred and direction of motion and its opposite:

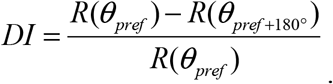

Contrast responses were fit using the Naka-Rushton equation (Naka and Rushton, 1966; Albrecht and Hamilton 1982; Heimel et al 2005):

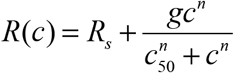

where *R*_*S*_ is the spontaneous firing rate, *c* is contrast, fitting parameters are contrast at half peak response (*c*_*50*_), gain (*g*), and exponent (*n*). Relative maximum gain (RMG) was calculated from the fits as maximum slope of the contrast response curve when the difference between maximum firing rate and spontaneous firing rate is normalized to 1. RMG indicates linearity of the contrast response curve, with 1 being completely linear. The contrast saturation index SI (Peirce, 2007) was defined to be

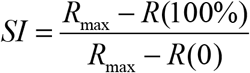

and indicates the degree of “supersaturation” of the contrast response (that is, the amount the response might be reduced at 100% contrast compared to the contrast that produces the maximum response, which might or might not be 100%).

Size tuning responses were fit to a product of two functions that represented the center response and the modulation of the center response, respectively:

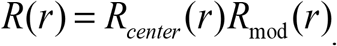

The center response *R*_*center*_ is the response of a Gaussian receptive field with the stimulus *S(x,y)*:

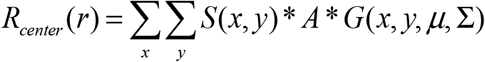

where *A* is the amplitude of the response (in spikes/sec), μ is the center position of the stimulus on the screen, and ∑ is the covariance matrix. Here, we constrained the Gaussian function to be circular by defining 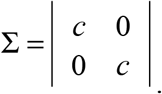.

The modulating function *R*_*mod*_*(r)* takes values between 0 and 2, and is proportional to the overlap of the stimulus and a circle of radius *R*_*max_stim*_:

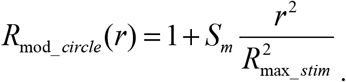

When the stimulus was an aperture, the modulating response was:

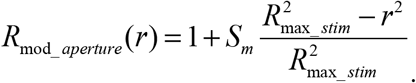

We quantified size tuning using two measurements, the size modulation parameter *Sm*, and the stimulus size when *R*_*center*_ exhibited half of its maximum response. *Sm* takes positive values for cells that exhibit length summation, and negative values for cells that exhibit surround suppression.

Cross orientation suppression was characterized in all cells using circular plaid stimuli consisting of two superimposed drifting sinusoidal gratings. One of the component gratings was assigned the previously established preferred orientation of the cell, the other component grating was assigned the orthogonal orientation. The direction of drift for the orthogonal grating was taken to be the preferred direction plus 90°. Response to the plaid stimulus can be related to the linear combination of responses to the component gratings:

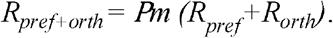

The plaid multiplier, *Pm*, is used to quantify the magnitude of the suppression of the response to the preferred orientation by the orthogonal orientation. Higher values of the plaid multiplier denote lower values of cross-orientation suppression.

### Computer code

All computer code related to this study is available at http://github.com/VH-lab/vhlab_publishedstudies and depends on libraries available at http://code.vhlab.org.

## Results

Our primary goal was to examine the influence of visual experience and age on the development of several receptive field properties. In particular, we were interested in uncovering whether cross-orientation suppression and size tuning depend on visual experience. To this end, we raised ferrets under three different conditions that were each designed to test a possible relationship between experience and selectivity. Typically-reared animals received 12 hours of visual experience daily, so they were exposed to complex natural scenes with objects of different sizes and mixtures of stimulus orientations. We recorded from animals living under typical rearing conditions at three different ages: postnatal day 40 (P40), postnatal day 60 (P60), and postnatal day 90 (P90). Dark-reared animals were raised in 24-hour constant darkness that was interrupted by two brief testing sessions (see Methods), and had very impoverished visual experience. A third group of animals was dark-reared but provided with two daily 1.5-hour “training” sessions for 5 days, where the animals were exposed to stimulation with sinusoidal gratings of a single orientation that occupied the full screen. Experience could be important for the development of cross-orientation-suppression or size tuning – furthermore experience with multiple orientations or objects of varying size might be necessary. The third experimental group was designed to test just that - whether the quality of visual experience influences the emergence of cross-orientation-suppression and size tuning.

We examined the refinement of receptive field properties at times related to the anatomical maturation of the visual cortical circuit (White and Fitzpatrick, 2007). Long-range horizontal connections extending for millimeters across the cortical surface exhibit adult-levels of complexity at around postnatal day 35-45 (Durack and Katz, 1996; Ruthazer and Stryker, 1996; White et al., 2001); volumetric synaptic density achieves adult levels around postnatal day 90 (Erisir and Harris, 2003). In principle, either of these anatomical features could underlie the development of cross-orientation suppression or size tuning. This study allowed us to connect these landmark events in the development of the visual cortical circuit to the changes in response properties of V1 neurons. A diagram of all animal groups is shown in **Figure 1.**

After isolating a cell, we assessed its direction tuning, followed by its spatial frequency tuning and temporal frequency tuning. All subsequent measurements were made with gratings that were aligned to the optimal direction, spatial frequency, and temporal frequency properties of the cell of interest. Then, we examined contrast responses, cross-orientation suppression, and size tuning. To our knowledge, the experience and age dependence of cross-orientation suppression and size tuning have not been examined previously, so we turn our attention to these results first, followed by orientation selectivity, direction selectivity, spatial and temporal frequency tuning, contrast responses, and evoked firing rates.

### Cross-orientation suppression

Cross-orientation suppression was robustly present in all animal groups in our study. We assessed cross-orientation suppression at several contrasts with a stimulus that was 10° by 10° in size. **Figure 2** shows responses from typical cells from animals in each experimental group. Responses to plaid drifting gratings were nearly always smaller than the sum of the individual responses to the two component directions. One of the two component gratings always had the cell’s preferred orientation and drifted in the preferred direction, while the other component grating had the orthogonal orientation (**Figure 3**).

**Figure 2.**
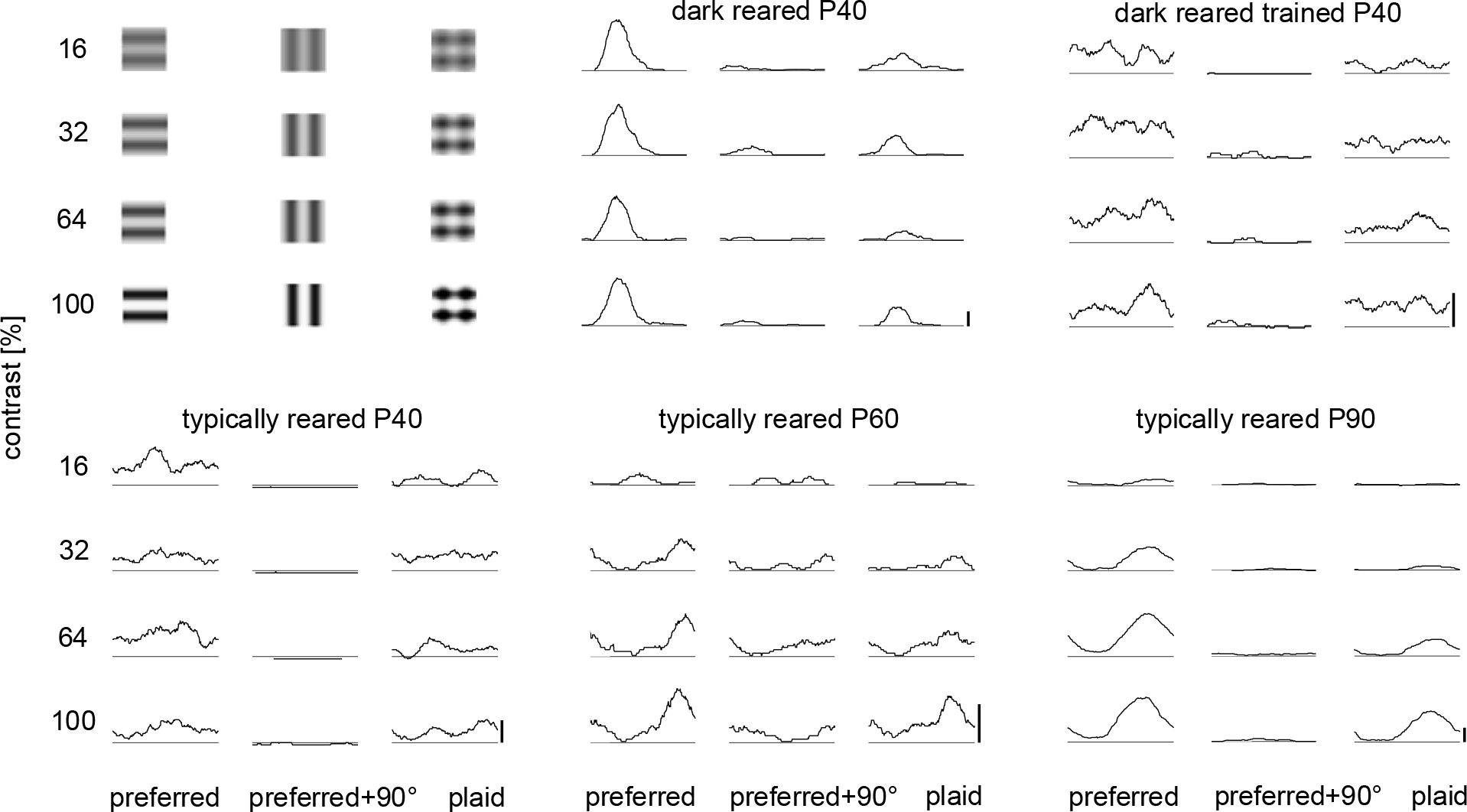
Representative cells in all experimental conditions exhibit cross-orientation suppression. For each cell, cycle-averaged responses to stimulation at the preferred direction, the orthogonal direction, and a plaid combination of the two stimuli are shown, for a variety of stimulus contrasts. Time is on the horizontal axis (showing 1 grating cycle). Vertical axis indicates response; bars denote 10 spikes/sec.

**Figure 3.**
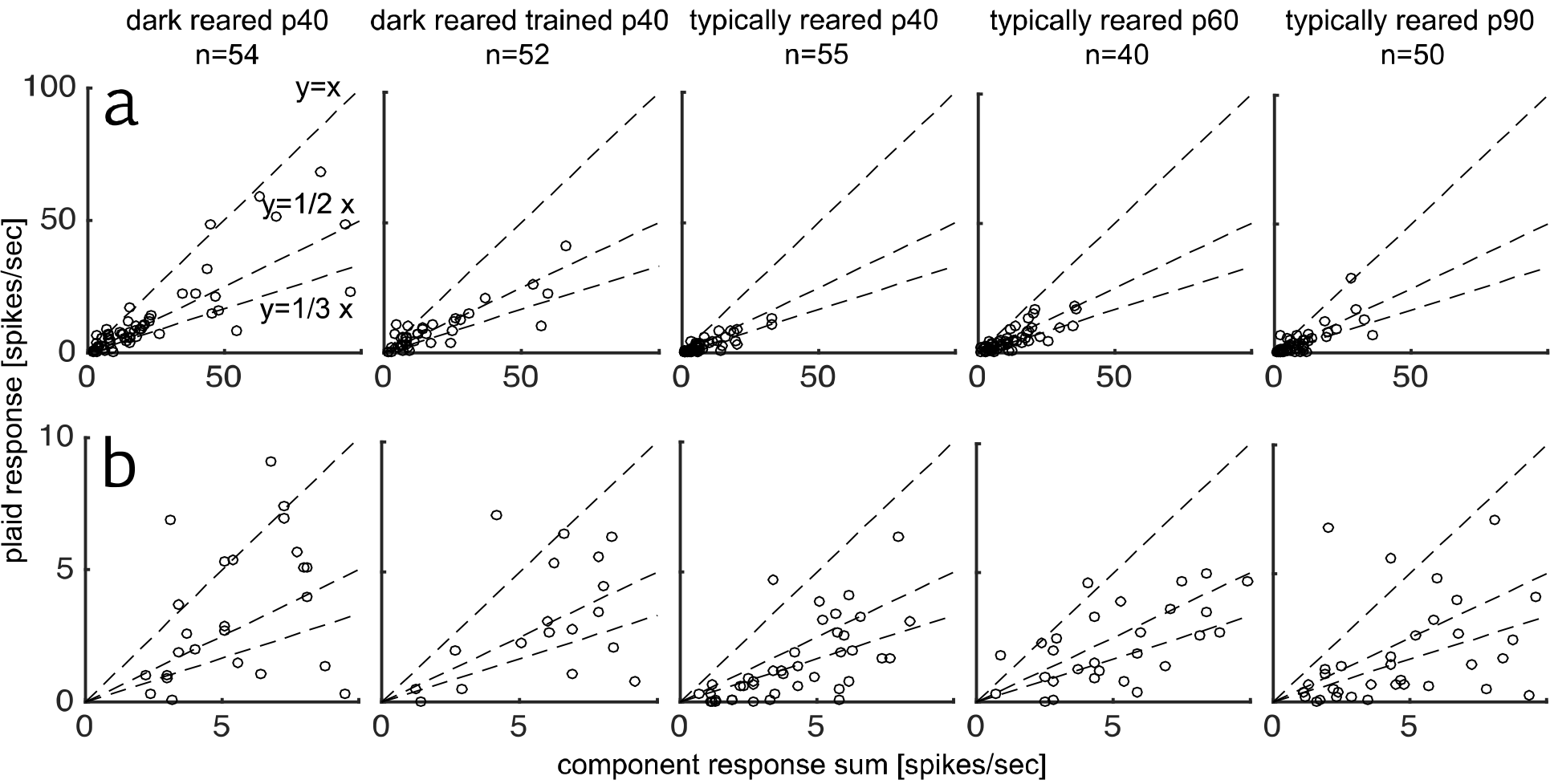
Cells in all animal groups exhibit robust cross-orientation suppression. Scatterplots of responses to plaid stimulation plotted against the linear sum of the response to preferred stimulation and orthogonal stimulation (that is, the component response sum) for stimuli of 100% contrast. Dashed lines show slopes y = x, y = 1/2 x, and y = 1/3 x. **a)** displays full range of data, **b)** displays data in the 0-10 spikes/sec range.

Although cross-orientation suppression was present in all experimental groups, the amount of suppression that we observed varied slightly with age and experience. We quantified cross-orientation suppression with a plaid multiplier *P*_*m*_, that compared the actual response to a plaid stimulus to the linear sum of the two components (**Figure 4**). A value of 1 would indicate perfect linear summation, and a value less than 1 indicates cross-orientation suppression. The Kruskal-Wallis H test shows a statistically significant effect of experimental condition on the plaid multiplier for all three contrast levels tested (32%: (χ^2^(4) = 22.15, *p < 0.005*; 64%: χ^2^(4) = 22.23, *p < 0.05*; 100%: χ^2^(4) = 29.09, *p < 0.05*). Interestingly, cross-orientation suppression in typically-reared animals showed an increase with experience at P40 (compared to dark-reared animals), but this initial increase in suppression was followed by a subsequent decrease at P60 and P90.

**Figure 4.**
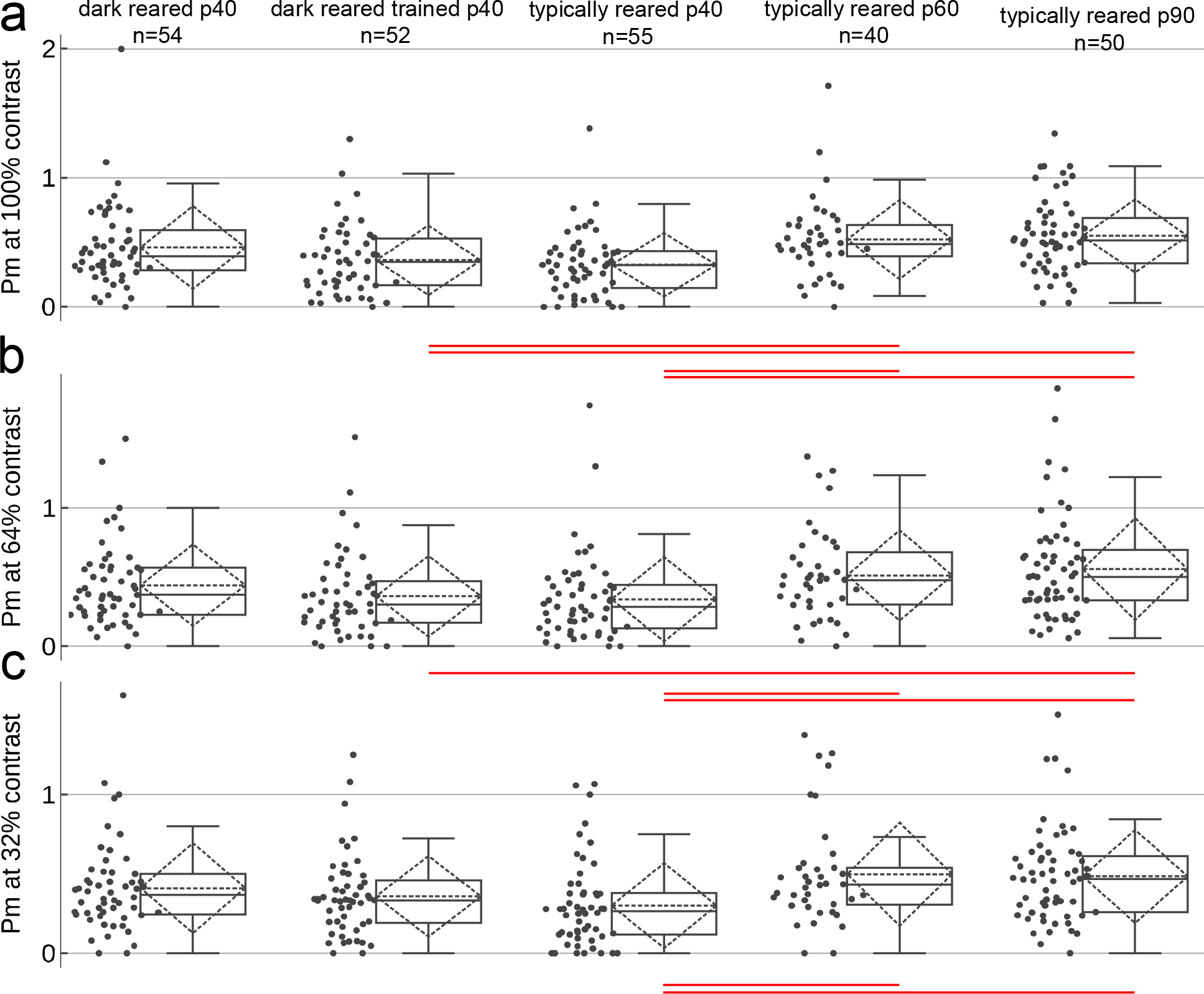
Median cross-orientation suppression varies slightly across age and condition. Plaid multiplier *P*_*m*_ is shown for all cells across all groups at 32%, 64%, and 100% contrast. For each group, all data points are shown at the left, and a box and whisker plot is shown at the right. The horizontal line in the center of each dotted diamond indicates the mean, and the dotted diamond tips are at ± 1 standard deviation. Lower *P*_*m*_ values indicate higher suppression. There is a slight increase in suppression in typically-reared P40 animals at all contrasts, and cross-orientation suppression reduces slightly at P60 and P90. Dashed lines are at 0.5 and 1. Red lines indicate pairwise differences significant at p<.05 (Bonferroni-corrected Kruskal-Wallis test).

### Size tuning

Size tuning is another parameter whose developmental profile has not been examined previously. Studies (Gilbert, 1977; Chisum et al., 2003) have shown that cells can exhibit a wide variety of responses to large stimuli. Some cells merely plateau in response to increasing stimulus size (**Figure 5a**). Other cells exhibit surround suppression, where stimulation outside of the classical receptive field induces weaker responses to central stimulation (**Figure 5b**). Finally, still other cells show length summation, where cells’ responses to stimuli that exceed the classical receptive field keep increasing (**Figure 5c**). We observed all of these response types in our animals.

**Figure 5.**
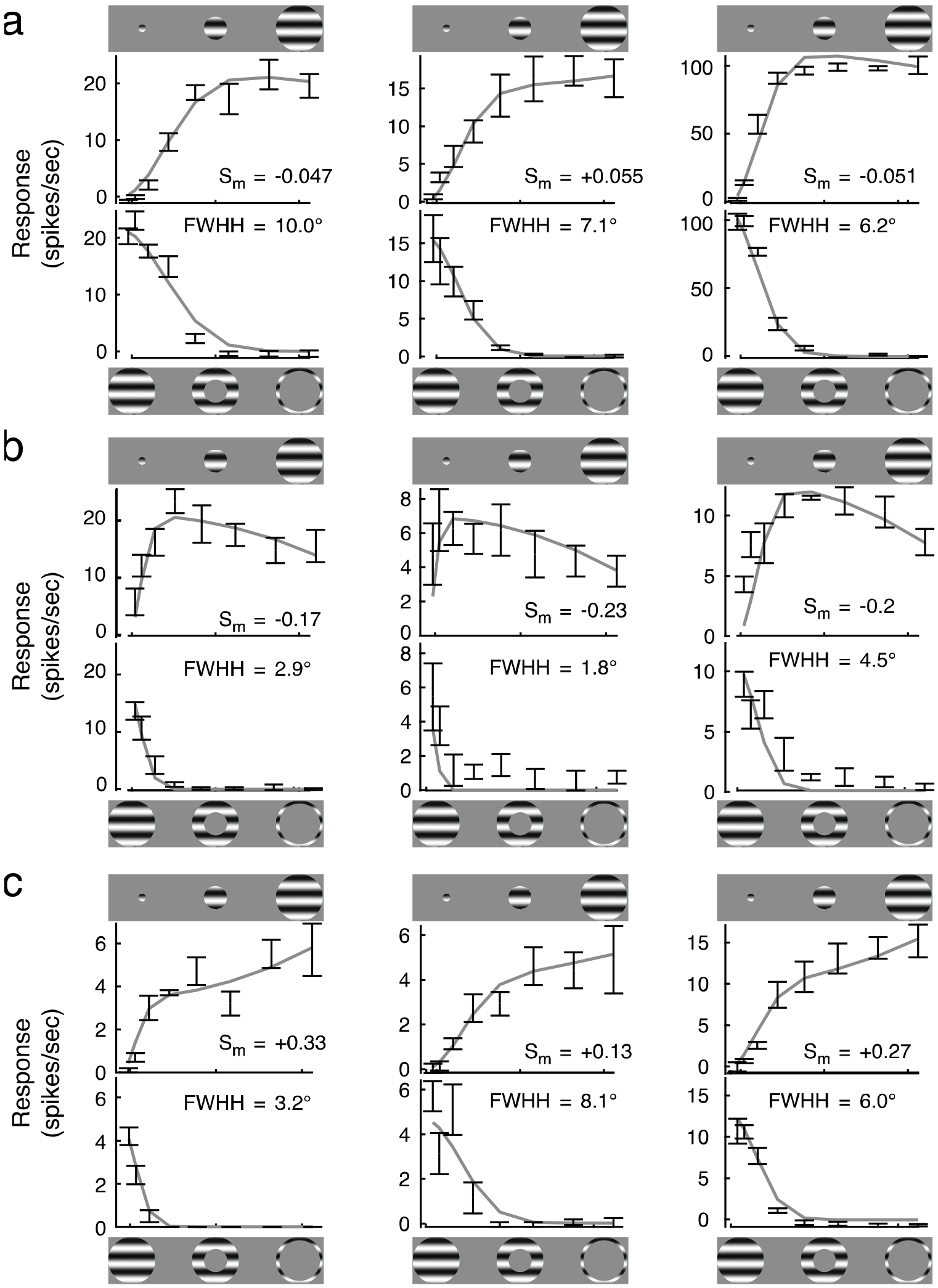
Diversity of size tuning curves in ferret visual cortex. Cells were fit to a Gaussian center and a circular modulating region (see Methods). Size tuning curves and corresponding fits for three broad groups of response categories: a) cells that exhibit a plateau but are not modulated by stimulation in the surround (*S*_*m*_ approximately 0), b) cells that exhibit surround suppression (*S*_*m*_ negative), and c) cells that exhibit length summation (*S*_*m*_ positive). Top panel for each cell shows responses to stimuli of increasing size [3°, 6°, 11°, 19°, 29°, 39°, 48°, 54°] while bottom panel shows responses to an annulus of increasing inner diameter (same sizes).Responses to both classes of stimuli were used to construct the fits. FWHH is full width at half height of the Gaussian center region.

To evaluate these responses quantitatively, we developed a fit function that included a Gaussian center component that was modulated by a circular surround component (see Methods). The degree of surround modulation was quantified by a single parameter, *Sm*, that was negative when stimulation of the surround was suppressive and was positive when the stimulation of the surround was enhancing. This parameter took values near 0 when surround stimulation did not influence the firing of the neuron. We examined responses to stimuli of increasing size and annular stimuli of decreasing inner diameter, and both sets of responses were used to establish the fits. Analysis of the responses to the annular stimulus allowed us to delineate the classical receptive field center (where stimulation evokes a response) from the surround that merely modulates responses to the center stimulus.

We observed plateau responses and surround suppression in all animal groups (**Figure 6a**), indicating that these variants of size tuning do not require visual experience for their expression. The fraction of neurons that exhibited substantial length summation increased in the oldest animals in the study (**Figure 6a**). Because we did not modulate visual experience in the oldest animals in our study, we cannot conclude whether these results are due to age or experience, but these responses do emerge at a time when the horizontal connections across the cortical surface have already reached anatomical maturity (Durack and Katz, 1996; Ruthazer and Stryker, 1996; White et al., 2001).

**Figure 6.**
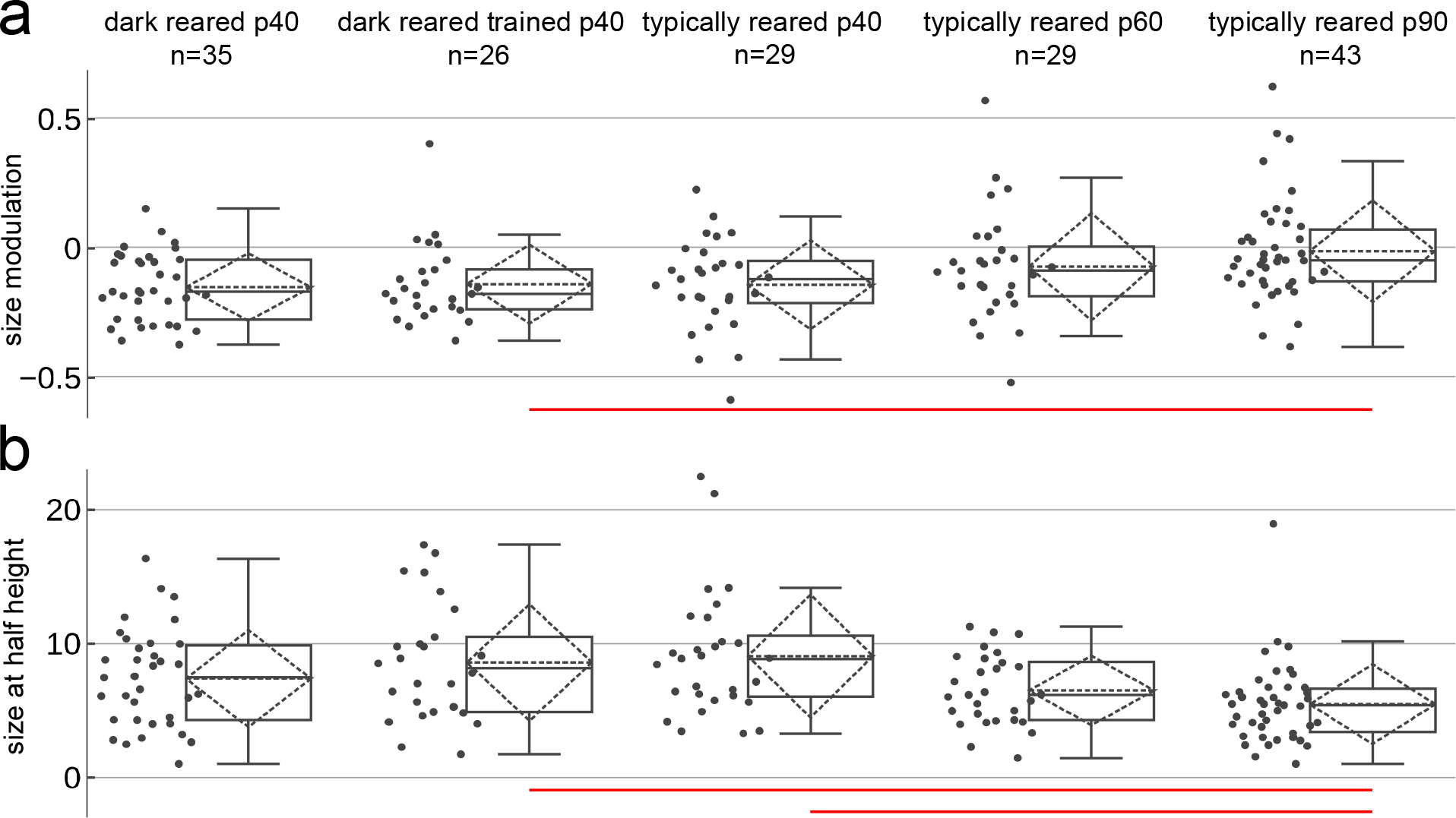
Size tuning properties with age and experience. a) Parameter *Sm*; cells that exhibit plateau responses and surround suppression are found in all animals, but cells that exhibited substantial length-summation were found more commonly in the oldest animals. Red lines indicate pairwise differences (Kruskal-Wallis test, Bonferroni corrected) at p<0.05). b) As expected from earlier reports, receptive field size as assessed by the full width at half height (FWHH) of the Gaussian center component exhibited decreases with experience, and was smallest in the oldest animals examined. Red lines indicate pairwise differences (paired t-test with Bonferroni correction) significant at p<0.05. Note that the major reduction of receptive field size occurs after other properties like orientation and direction selectivity have been established.

Consistent with prior reports, we observed a substantial decrease of receptive field center size in the oldest animals (**Figure 6b**). Median receptive field center sizes dropped from 7.5° in dark-reared animals to 5.4° in the P90 animals. This maturation coincides with the peak synapse density in layer 2/3 in ferret (around P90) (Erisir and Harris, 2003; White and Fitzpatrick, 2007).

### Orientation and direction selectivity

Previous studies have found that orientation selectivity is present at the time of eye opening and that it increases with the onset of visual experience (Chapman and Stryker, 1993; White et al., 2001; Li et al., 2006). Direction selectivity, on the other hand, is almost entirely absent at the time of eye opening, emerges over several days, and requires visual experience (Li et al., 2006). Our results largely recapitulated these prior observations: dark-reared animals exhibited moderately strong orientation selectivity that increased with visual experience, and dark-reared animals exhibited very weak direction selectivity that was also increased by visual experience (**Figure 7**).

**Figure 7.**
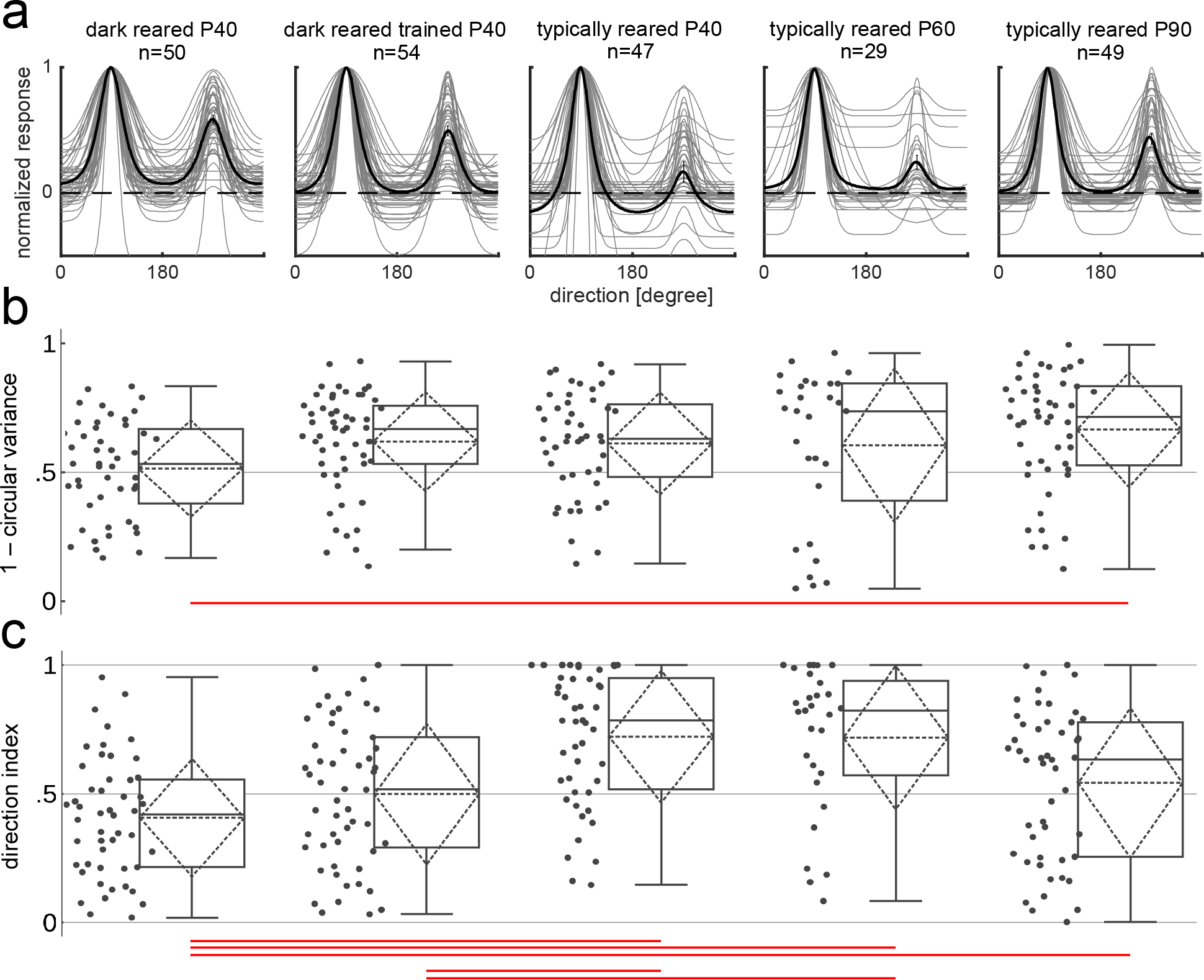
Effects of experimental condition on the development of orientation and direction tuning. a) Normalized direction tuning curves for the 5 experimental conditions; b) orientation selectivity quantified as 1-circular variance; c) direction selectivity quantified as direction index. Dashed lines are at 0.5 DI and 1-circular variance; red lines indicate pairwise differences significant at p<0.05 (Kruskal-Wallis test, Bonferroni correction). As expected from previous studies, both orientation selectivity and direction selectivity exhibit increases with experience. Unexpectedly, direction selectivity reached a peak at P40/P60 and reduced slightly at P90, consistent with the idea that direction selectivity does not develop in a monotonic manner.

These data also give us a new view of the impact of visual experience with simple grating stimuli on orientation and direction selectivity. Previous work has demonstrated that only a few hours (3-9 hours) of experience with a moving visual stimulus is sufficient to cause the rapid emergence of direction selectivity and a concurrent increase in orientation selectivity (Li et al., 2008; Van Hooser et al., 2012; Ritter et al., 2017). However, for methodological reasons, these experiments only assessed orientation and direction selectivity immediately after exposure to the “training stimulus”, leaving open the possibility that the effects of such visual experience were transient. In this study, recordings were obtained 1-5 days after the last training session, allowing us to address this possibility. Dark-reared and trained animals exhibited direction selectivity that was intermediate between dark-reared animals that did not have training and animals with typical visual experience. P40 dark-reared animals exhibited direction selectivity that was significantly lower than P40 typically-reared animals. A direct comparison between dark-reared animals and trained dark-reared animals did not reach significance (p=0.09, Kruskal-Wallis test), but trained and dark-reared animals exhibited direction selectivity that was not different from typically-reared P90 animals (**Figure 7c**; p<0.05, Kruskal-Wallis test, Bonferroni correction), which show some substantial direction selectivity. This evidence suggests that the exposure to the training stimulus did produce relatively lasting, if small, changes in receptive field properties.

Finally, we were surprised to observe that direction selectivity changes non-monotonically with age. We observed the strongest direction selectivity in P40 animals that were typically reared. Direction selectivity index values decreased at P90, but still remained well above the values of visually-naïve animals. This result suggests that selectivity for some features goes through periods of increases and decreases as the animal matures.

### Spatial and temporal frequency tuning

Spatial frequency preference showed a substantial and expected dependency on experience. Spatial frequency was characterized in all cells using drifting sinusoidal grating stimuli of varying spatial frequency ([0.05, 0.1, 0.15, 0.2, 0.3, 0.5, 0.8] cycles per degree visual angle), 4Hz temporal frequency, 100% contrast, and orientation and direction fixed at the previously established optimal value for each cell. As expected from previous reports, cells from animals in the P40 age group prefer lower spatial frequencies regardless of rearing condition (**Figure 8ab**), consistent with the lower resolution of vision in younger animals (Freeman and Marg, 1975; DeAngelis et al., 1993; Tavazoie and Reid, 2000; Heimel et al., 2007). In the two older groups, P60 and P90, spatial frequency preference shifted towards higher frequencies. There was also a noticeable diversification of SF preferences with age – SF preferences of individual cells in younger animals were more tightly clustered. Conversely, in older animals, cells preferred higher SFs, and their preferences had a larger spread.

**Figure 8.**
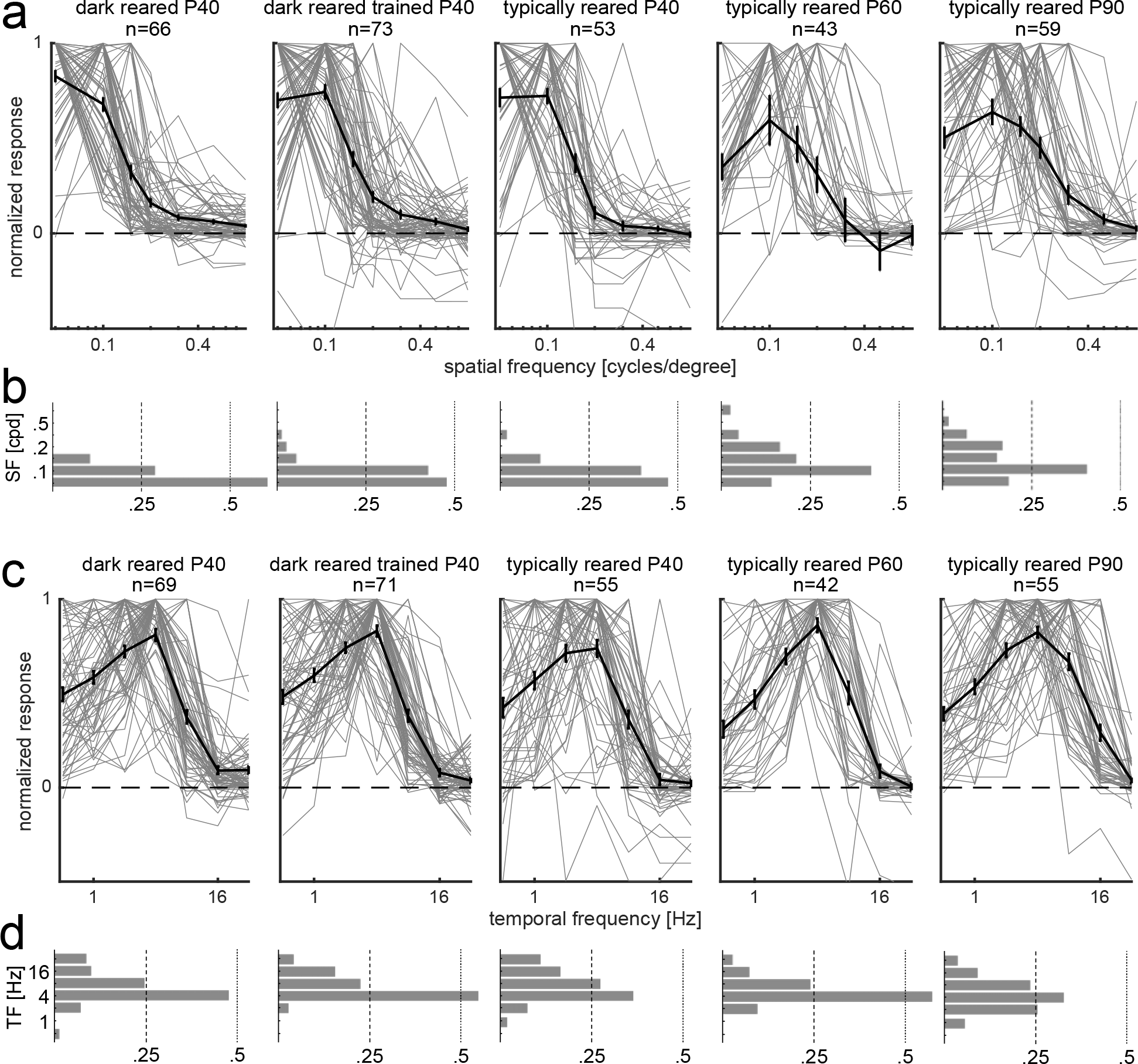
Effects of experimental condition on the development of spatial and temporal frequency preference. a) Normalized spatial frequency tuning curves for the 5 experimental conditions; b) normalized spatial frequency histogram; ticks on the y axis denote 0.05, 0.1, 0.15, 0.2, 0.3, 0.5, and 0.8 cycles per degree (cpd) (bottom to top). c) Normalized temporal frequency tuning curves for the 5 experimental conditions; d) normalized temporal frequency histogram; ticks on the y axis denote 0.5, 1, 2, 4, 8, 16, and 32 Hz (bottom to top). As expected from previous studies, and consistent with decreases in receptive field size that are reported in **Figure 6**, spatial frequency preferences exhibited slight increases with age, indicating that the spatial resolution of visual processing increases with age. Temporal frequency preferences were relatively constant over the ages and rearing conditions studied here.

Temporal frequency preferences were subject to fewer differences across the experimental groups compared to selectivity to other features. Temporal frequency was characterized in all cells using drifting sinusoidal grating stimuli of varying temporal frequency ([0.5, 1, 2, 4, 8, 16, 32] Hz), 100% contrast, and spatial frequency, orientation, and direction fixed at the previously established optimal value for each cell. Cells from animals in the P40 and P60 age groups preferred lower temporal frequencies regardless of rearing condition (**Figure 8cd**). In the P90 age group, TF preference shifted slightly towards higher frequencies, but these changes were quite modest.

### Contrast tuning

Contrast tuning was subject to subtle differences across the different animal groups. Contrast responses were examined in all cells using drifting sinusoidal grating stimuli of varying contrast (2%, 4%, 8%, 16%, 32%, 64%, 100%), and with temporal frequency, spatial frequency, and direction fixed at the previously established optimal values for each cell (**Figure 9**). There were no statistically significant differences among relative maximum gain (RMG) for different conditions (**Figure 9b**). There were small but significant differences in the amount of “supersaturation” that was exhibited by neurons in this different groups. The median value for all groups was very close to 0 (no supersaturation) but a few cells were substantially suppressed at the highest contrast (**Figure 9c**). Overall, visual experience and age appeared to have only a modest impact on contrast responses.

**Figure 9.**
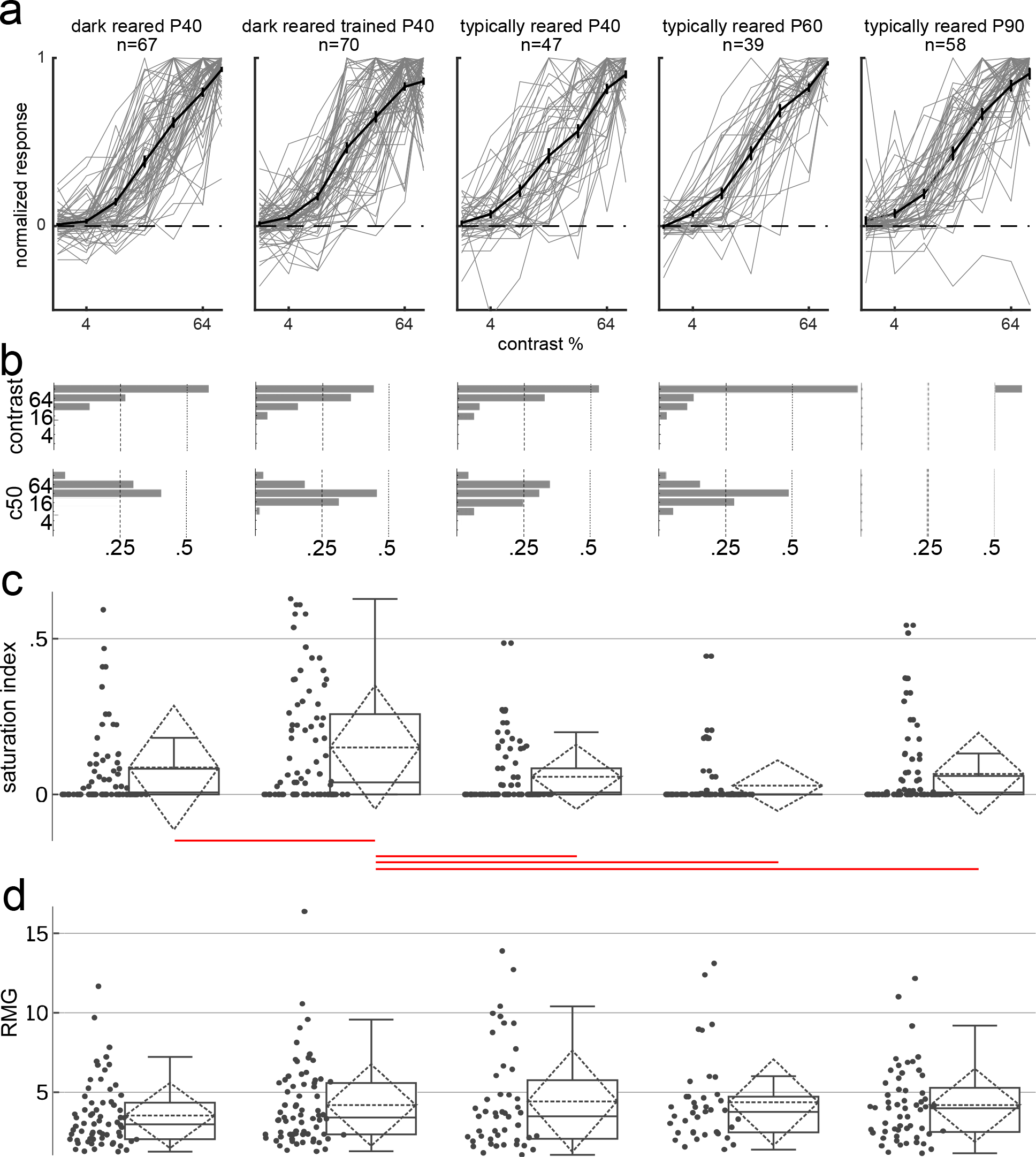
Effects of experimental condition on the development of contrast preference. a) Normalized contrast tuning curves for the 5 experimental conditions; b) contrast at peak (top), and half-peak response (bottom); c) saturation index of contrast response curve; d) linearity in response to contrast quantified by relative maximum gain, lower values indicate more linearity. Red lines indicate pairwise differences significant at p<0.05. Contrast responses were relatively constant over the developmental ages and rearing conditions studied here, with some slight variation in saturation index.

### Firing rate

One major parameter that exhibited a large change with age was evoked firing rate. The maximum evoked firing rate was taken to be the strongest trial-averaged response to sinusoidal grating stimulation that we observed – that is, the response measured at the preferred direction, spatial frequency, temporal frequency, and best contrast. Evoked maximum firing rates began at around 10Hz in young animals, but increased substantially in the P90 animals to about 20Hz on average (**Figure 10**). Kruskal-Wallis H test shows a statistically significant effect of experimental condition on firing rate (χ^2^(4) = 23.00, *p < 0.05*). We observed no difference in median evoked firing rate across the three rearing conditions at P40, but large firing rates (>40Hz) were only observed in the two older groups. These findings suggest that the network changes that support high firing rates are still emerging, even after 1 month of visual experience (P60).

**Figure 10.**
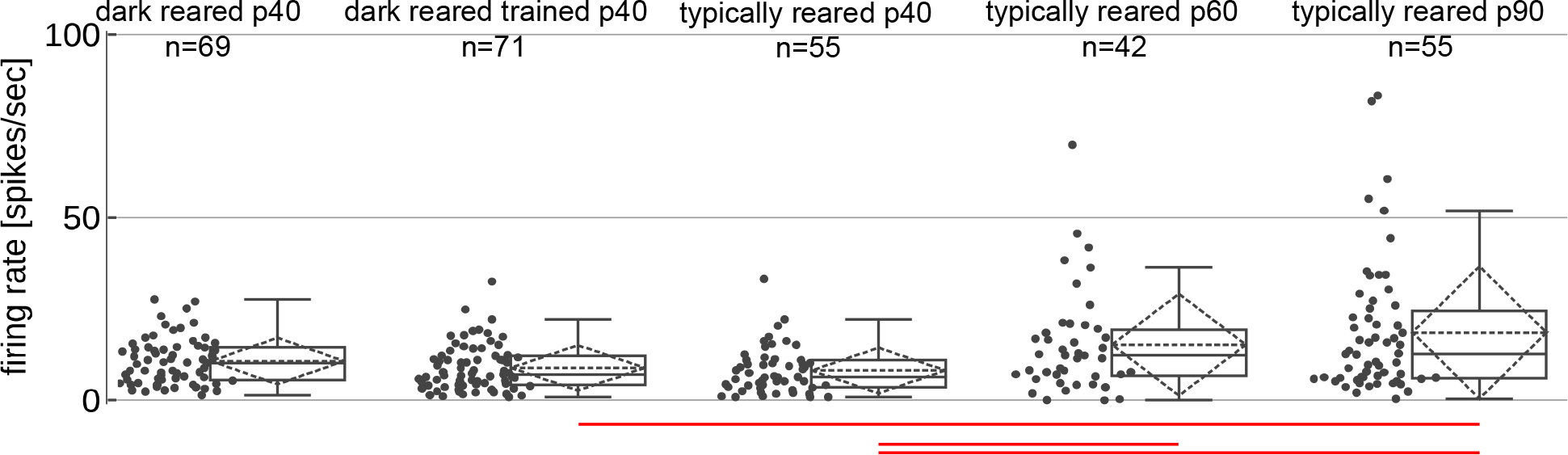
Firing rate across experimental conditions. Red lines indicate pairwise differences (Kruskal Wallis test, Bonferroni corrected) significant at p<0.05. The oldest animals exhibited substantially higher firing rates than younger animals.

## Discussion

We characterized the role of visual experience and age on the development of V1 receptive field properties in ferret. We found that cross-orientation suppression and surround suppression are present regardless of whether the animal has had any experience with visual stimuli. In addition, we found that increases in direction selectivity that are produced by short-term exposures to moving visual stimuli are retained over days. Direction selectivity reached a peak at P40 and decreased slightly in older animals. Finally, we observed that two features, length summation and high evoked firing rates, were primarily found in the oldest animals (P90).

### Contribution of sensory experience to normalization

Normalization and contextual interactions, including cross-orientation suppression and size tuning, are canonical computations of neural circuits (Carandini and Heeger, 2011; Angelucci et al., 2017). Cross-orientation suppression is a form of response normalization that has been observed in a wide variety of cortical areas, including V1 (Gizzi et al., 1990; Ringach et al., 2002), V2 (Rowekamp and Sharpee, 2017), V4 (Reynolds et al., 1999), MT (Britten and Heuer, 1999; Heuer and Britten, 2002), and IT (Zoccolan et al., 2005). It is ubiquitous across examined species (Carandini and Heeger, 2011), being found in macaque (Ringach et al., 2002), cat (DeAngelis et al., 1992), mouse (Sato et al., 2016), and even *Drosophila* (Olsen et al., 2010). Brain imaging studies have also found that contrast-dependent suppression is precisely maintained across the entire neural population (Busse et al., 2009; MacEvoy et al., 2009). Surround suppression is also ubiquitous across species, being found in mouse (Van den Bergh et al., 2010; Self et al., 2014), ferret (Rubin et al., 2015), cat (Hubel and Wiesel, 1965; Blakemore and Tobin, 1972; Gilbert, 1977; DeAngelis et al., 1994; Sengpiel et al., 1997), monkey (Hubel and Wiesel, 1968; Sceniak et al., 1999; Jones et al., 2001; Cavanaugh et al., 2002a, b), and human (Williams et al., 2003; Zenger-Landolt and Heeger, 2003).

Due to the importance of normalization and contextual interactions to selectivity in the presence of multiple stimuli, including in natural scenes (Barlow, 1961, 1972; Bauman and Bonds, 1991; Somers et al., 1995; Carandini and Ringach, 1997; Lauritzen et al., 2001; Vinje and Gallant, 2002; David et al., 2004; Nurminen and Angelucci, 2014; Angelucci et al., 2017), it seemed possible that sensory experience with multiple stimuli or objects of various sizes might be necessary for the expression of cross-orientation suppression and surround suppression, or at least its refinement. Our results provide strong evidence that sensory experience is unnecessary for the development of cross-orientation suppression or surround suppression. Like orientation selectivity, these features are robustly present at the onset of visual experience and in dark-reared animals, and were not greatly impacted by varying levels of experience.

### Cortical circuits and normalization

The circuit mechanisms of cross-orientation suppression and size tuning are unknown. Some models posit that local connections within the cortex provide suppression, either via increased inhibition or reduced excitation (Somers et al., 1995; Haider et al., 2010; Sato et al., 2014; Rubin et al., 2015). Other models suggest that cross-orientation suppression reflects reduced synchronized input from lateral geniculate nucleus when multiple stimuli are present (Priebe and Ferster, 2006), or that the inhibition arises via feedback connections from higher cortical areas (Angelucci et al., 2002; Angelucci et al., 2017). Our results do not allow us to choose among these alternatives, but we can make one inference. Dark-reared animals showed strong cross-orientation suppression and typical rates of surround suppression, and prior research has noted that the long-range horizontal connections across the cortical surface are poorly formed in dark-reared animals (White et al., 2001). Therefore, it is unlikely that the long-range horizontal connections are a critical component for cross-orientation or surround suppression.

We observed the highest percentage of length-summing cells and the highest evoked firing rates in animals that had attained approximately 2 months of visual experience (P90). What circuit properties are modified at this time? The long-range horizontal connections are anatomically established by P35-45 (Durack and Katz, 1996; Ruthazer and Stryker, 1996; White et al., 2001), but excitatory synaptic density in layer 2/3 does not reach its maximum until P80-100 (Erisir and Harris, 2003). Further, feedback connections to V1 from extrastriate areas are present at the time of eye opening, and are pruned from eye opening to P70 (Khalil and Levitt, 2014). The refinement of these circuit elements could contribute to the emergence of normalization related tuning properties.

### Influence of experience on direction selectivity

The development of direction selectivity requires visual experience. Visually-naïve animals do not exhibit strong direction selectivity, and animals that are dark reared throughout an early critical period do not attain direction selectivity even if they are subsequently exposed to light for weeks (Li et al., 2006). Previous experiments have found that just 3-9 hours of visual experience with moving stimuli is sufficient to cause emergence of direction selectivity in visually-naïve, anesthetized ferrets (Li et al., 2008; Van Hooser et al., 2012; Roy et al., 2016; Ritter et al., 2017). Previous experiments were done in acute preparations, and it was unknown if these effects would persist for more than a few hours. We exposed dark-reared animals to 14-16 hours of stimulation with large drifting grating stimuli. Stimulus delivery differed from previous studies in which the anesthetized ferrets were paralyzed to eliminate eye movements. Here, awake animals were head-fixed but were free to move their eyes. Due to their age, the animals often spent tens of minutes of their 80 minute exposures sleeping. Further, trained dark-reared animals spent the vast majority of their total time after eye opening in darkness, which might have degraded any experience-dependent changes.

Nevertheless, we found that the P40 animals that were dark reared and exposed to large moving gratings exhibited average direction selectivity with a magnitude between those of P40 dark-reared animals and P40 typically-reared animals. While the differences in direction selectivity between P40 dark-reared and dark-reared trained animals did not reach significance, P40 dark-reared and trained animals exhibited direction selectivity that was not different from P90 animals, which is evidence of modestly increased selectivity. This is evidence of the persistence of the influence of visual experience on direction selectivity.

We also found a surprising decline of direction selectivity in the second month of visual experience (approximately P60-P90). Direction selectivity index values peaked at P40, and declined slightly afterwards. This suggests that the changes to direction selectivity over age are non-monotonic, with direction selectivity increasing and decreasing as visual circuitry reaches maturity.

### Role of experience in development of primary sensory receptive field properties

One could imagine two broad ideas about how receptive field properties might be formed in primary sensory areas. Visual circuits could analyze the input statistics and design appropriate filters to encode this information. Indeed, when artificial cortical networks with learning rules are presented with natural scenes, the early filters in these networks resemble those of visual cortical neurons (Olshausen and Field, 1996; Bell and Sejnowski, 1997; van Hateren and van der Schaaf, 1998; Ranzato et al., 2007). Thus, in principle, it is possible that these response properties could be derived purely from experience.

But an alternative hypothesis – one that posits that efficient receptive field properties have been genetically derived over eons of evolution – seems more consistent with experimental data. Orientation tuning, spatial and temporal frequency tuning, and normalization are present at the onset of the visual experience, and are only modestly altered by normal experience (Chapman and Stryker, 1993; DeAngelis et al., 1993; White et al., 2001; this paper; Li et al., 2006) (though abnormal experience can create highly aberrant receptive field properties, as in Mitchell, 1988; White et al., 2001; Prusky and Douglas, 2003). Experience is necessary for development of cortical direction selectivity (Li et al., 2006), but the tuning parameters that will emerge (angle preference and speed tuning) are already determined, and experience seems only to enhance the selectivity (Li et al., 2008; Roy et al., 2016; Ritter et al., 2017). The major contribution of experience, at least in primary visual cortex, appears to be the alignment of the inputs of the two eyes (Wang et al., 2010), some refinement of spatial frequency preferences (Mitchell, 1988), and the establishment of appropriate gains for selectivity (Turrigiano and Nelson, 2000).

These results provide one of two possible conclusions. It could be the case that the major properties of V1 neurons (except ocular alignment) are simply established by experience-independent mechanisms, such as molecular cues and modifications due to spontaneous activity (Meister et al., 1991; Ruthazer and Stryker, 1996; Cang et al., 2008). Or, perhaps, there are important experience-dependent modifications, but these modifications are not observable with the stimuli we have used here, and would only be apparent when animals are viewing more natural stimuli (e.g., Berkes et al., 2011).

Experience is clearly important to the mammalian brain, but whether its influence is instructive may depend considerably on the brain area or, as suggested here, on the type of neural computation. The evidence presented here suggests that cross-orientation suppression and surround suppression are present in ferrets independent of visual experience.

## Acknowledgements

This work was funded by the NSF IOS 1120938 (SDV, JF,MP), and by a Hoffman Research Fellowship from Bates College (CEO). We thank Alexandra Hempel, Victoria Drumm, David Landesman, George Popa, Tudor Dragoi, Rebecca Panitch, and Lizbeth Lueck, for help with animal husbandry, and we thank members of the Van Hooser lab for comments.

## References

Adelson EH, Movshon JA (1982) Phenomenal coherence of moving visual patterns. Nature 300:523–525.

Albrecht DG, Hamilton DB (1982) Striate cortex of monkey and cat: contrast response function. J Neurophysiol 48:217–237.

Angelucci A, Levitt JB, Lund JS (2002) Anatomical origins of the classical receptive field and modulatory surround field of single neurons in macaque visual cortical area V1. Prog Brain Res 136:373–388.

Angelucci A, Bijanzadeh M, Nurminen L, Federer F, Merlin S, Bressloff PC (2017) Circuits and Mechanisms for Surround Modulation in Visual Cortex. Annu Rev Neurosci.

Barlow HB (1961) Possible principles underlying the transformation of sensory messages. In: Sensory Communication, pp 217–234. Cambridge, MA: MIT Press.

Barlow HB (1972) Single units and sensation: a neuron doctrine for perceptual psychology? Perception 1:371–394.

Batschelet E (1981) Circular statistics in Biology. New York: Academic Press.

Bauman LA, Bonds AB (1991) Inhibitory refinement of spatial frequency selectivity in single cells of the cat striate cortex. Vis Res 31:933–944.

Bell AJ, Sejnowski TJ (1997) The “independent components” of natural scenes are edge filters. Vision Res 37:3327–3338.

Berkes P, Orban G, Lengyel M, Fiser J (2011) Spontaneous cortical activity reveals hallmarks of an optimal internal model of the environment. Science 331:83–87.

Blakemore C, Tobin EA (1972) Lateral inhibition between orientation detectors in the cat's visual cortex. Exp Brain Res 15:439–440.

Bolz J, Gilbert CD (1986) Generation of end-inhibition in the visual cortex via interlaminar connections. Nature 320:362–365.

Bolz J, Gilbert CD (1989) The Role of Horizontal Connections in Generating Long Receptive Fields in the Cat Visual Cortex. Eur J Neurosci 1:263–268.

Britten KH, Heuer HW (1999) Spatial summation in the receptive fields of MT neurons. J Neurosci 19:5074–5084.

Busse L, Wade AR, Carandini M (2009) Representation of concurrent stimuli by population activity in visual cortex. Neuron 64:931–942.

Cang J, Niell CM, Liu X, Pfeiffenberger C, Feldheim DA, Stryker MP (2008) Selective disruption of one Cartesian axis of cortical maps and receptive fields by deficiency in ephrin-As and structured activity. Neuron 57:511–523.

Carandini M, Ringach DL (1997) Predictions of a recurrent model of orientation selectivity. Vision Res 37:3061–3071.

Carandini M, Heeger DJ (2011) Normalization as a canonical neural computation. Nat Rev Neurosci 13:51–62.

Carandini M, Heeger DJ, Movshon JA (1997) Linearity and Normalization in Simple Cells of the Macaque Primary visual Cortex. J Neurosci 17:8621–8644.

Cavanaugh JR, Bair W, Movshon JA (2002a) Selectivity and spatial distribution of signals from the receptive field surround in macaque V1 neurons. J Neurophysiol 88:2547–2556.

Cavanaugh JR, Bair W, Movshon JA (2002b) Nature and interaction of signals from the receptive field center and surround in macaque V1 neurons. J Neurophysiol 88:2530–2546.

Chapman B, Stryker MP (1993) Development of orientation selectivity in ferret visual cortex and effects of deprivation. J Neurosci 13:5251–5262.

Chisum HJ, Mooser F, Fitzpatrick D (2003) Emergent properties of layer 2/3 neurons reflect the collinear arrangement of horizontal connections in tree shrew visual cortex. J Neurosci 23:2947–2960.

Clemens JM, Ritter NJ, Roy A, Miller JM, Van Hooser SD (2012) The laminar development of direction selectivity in ferret visual cortex. J Neurosci 32:18177–18185.

David SV, Vinje WE, Gallant JL (2004) Natural stimulus statistics alter the receptive field structure of vl neurons. J Neurosci 24:6991–7006.

DeAngelis GC, Ohzawa I, Freeman RD (1993) Spatiotemporal organization of simple-cell receptive fields in the cat's striate cortex. I. General characteristics and postnatal development. J Neurophysiol 69:1091–1117.

DeAngelis GC, Freeman RD, Ohzawa I (1994) Length and width tuning of neurons in the cat's primary visual cortex. J Neurophysiol 71:347–374.

DeAngelis GC, Robson JG, Ohzawa I, Freeman RD (1992) Organization of suppression in receptive fields of neurons in cat visual cortex. J Neurophysiol 68:144–163.

Durack JC, Katz LC (1996) Development of horizontal projections in layer 2/3 of ferret visual cortex. Cereb Cortex 6:178–183.

Erisir A, Harris JL (2003) Decline of the critical period of visual plasticity is concurrent with the reduction of NR2B subunit of the synaptic NMDA receptor in layer 4. J Neurosci 23:5208–5218.

Freeman DN, Marg E (1975) Visual acuity development coincides with the sensitive period in kittens. Nature 254:614–615.

Gilbert CD (1977) Laminar differences in receptive field properties of cells in cat primary visual cortex. J Physiol 268:391–421.

Gizzi MS, Katz E, Schumer RA, Movshon JA (1990) Selectivity for orientation and direction of motion of single neurons in cat striate and extrastriate visual cortex. J Neurophysiol 63:1529–1543.

Haider B, Krause MR, Duque A, Yu Y, Touryan J, Mazer JA, McCormick DA (2010) Synaptic and network mechanisms of sparse and reliable visual cortical activity during nonclassical receptive field stimulation. Neuron 65:107–121.

Heeger DJ (1992) Normalization of cell responses in cat striate cortex. Vis Neurosci 9:181–198.

Heimel JA, Van Hooser SD, Nelson SB (2005) Laminar organization of response properties in primary visual cortex of the gray squirrel (Sciurus carolinensis). J Neurophysiol 94:3538–3554.

Heimel JA, Hartman RJ, Hermans JM, Levelt CN (2007) Screening mouse vision with intrinsic signal optical imaging. Eur J Neurosci 25:795–804.

Heuer HW, Britten KH (2002) Contrast dependence of response normalization in area MT of the rhesus macaque. J Neurophysiol 88:3398–3408.

Hubel DH, Wiesel TN (1965) Receptive Fields And Functional Architecture In Two Nonstriate Visual Areas (18 And 19) Of The Cat. J Neurophysiol 28:229–289.

Hubel DH, Wiesel TN (1968) Receptive fields and functional architecture of monkey striate cortex. J Physiol 195:215–243.

Jones HE, Grieve KL, Wang W, Sillito AM (2001) Surround suppression in primate VI. J Neurophysiol 86:2011–2028.

Khalil R, Levitt JB (2014) Developmental remodeling of corticocortical feedback circuits in ferret visual cortex. J Comp Neurol 522:3208–3228.

Lauritzen TZ, Krukowski AE, Miller KD (2001) Local correlation-based circuitry can account for responses to multi-grating stimuli in a model of cat V1. J Neurophysiol 86:1803–1815.

Li Y, Fitzpatrick D, White LE (2006) The development of direction selectivity in ferret visual cortex requires early visual experience. Nat Neurosci 9:676–681.

Li Y, Van Hooser SD, Mazurek M, White LE, Fitzpatrick D (2008) Experience with moving visual stimuli drives the early development of cortical direction selectivity. Nature 456:952–956.

MacEvoy SP, Tucker TR, Fitzpatrick D (2009) A precise form of divisive suppression supports population coding in the primary visual cortex. Nat Neurosci 12:637–645.

Mazurek M, Kager M, Van Hooser SD (2014) Robust quantification of orientation selectivity and direction selectivity. Front Neural Circuits 8:92.

Meister M, Wong ROL, Baylor DA, Shatz CJ (1991) Synchronous bursts of action-potentials in ganglion cells of the developing mammalian retina. Science 252:939–943.

Mitchell DE (1988) The extent of visual recovery from early monocular or binocular visual deprivation in kittens. J Physiol 395:639–660.

Morrone MC, Burr DC, Maffei L (1982) Functional implications of cross-orientation inhibition of cortical visual cells. I. Neurophysiological evidence. Proc R Soc Lond B 216:335–354.

Morrone MC, Burr DC, Speed HD (1987) Cross-orientation inhibition in cat is GABA mediated. Expl Brain Res 67:635–644.

Naka KI, Rushton WA (1966) S-potentials from colour units in the retina of fish (Cyprinidae). J Physiol 185:536–555.

Nurminen L, Angelucci A (2014) Multiple components of surround modulation in primary visual cortex: multiple neural circuits with multiple functions? Vision Res 104:47–56.

Ohshiro T, Angelaki DE, DeAngelis GC (2011) A normalization model of multisensory integration. Nat Neurosci 14:775–782.

Olsen SR, Bhandawat V, Wilson RI (2010) Divisive normalization in olfactory population codes. Neuron 66:287–299.

Olshausen BA, Field DJ (1996) Emergence of simple-cell receptive field properties by learning a sparse code for natural images. Nature 381:607–609.

Peirce JW (2007) The potential importance of saturating and supersaturating contrast response functions in visual cortex. J Vis 7:13.

Priebe NJ, Ferster D (2006) Mechanisms underlying cross-orientation suppression in cat visual cortex. Nat Neurosci 9:552–561.

Prusky GT, Douglas RM (2003) Developmental plasticity of mouse visual acuity. Eur J Neurosci 17:167–173.

Ranzato M, Boureau Y-L, LeCun Y (2007) Sparse feature learning for deep belief networks. In: NIPS.

Reynolds JH, Heeger DJ (2009) The normalization model of attention. Neuron 61:168–185.

Reynolds JH, Chelazzi L, Desimone R (1999) Competitive mechanisms subserve attention in macaque areas V2 and V4. The Journal of neuroscience: the official journal of the Society for Neuroscience 19:1736–1753.

Ringach DL, Bredfeldt CE, Shapley RM, Hawken MJ (2002) Suppression of neural responses to nonoptimal stimuli correlates with tuning selectivity in macaque V1. J Neurophysiol 87:1018–1027.

Ritter NJ, Anderson NM, Van Hooser SD (2017) Visual Stimulus Speed Does Not Influence the Rapid Emergence of Direction Selectivity in Ferret Visual Cortex. J Neurosci 37:1557–1567.

Rowekamp RJ, Sharpee TO (2017) Cross-orientation suppression in visual area V2. Nature communications 8:15739.

Roy A, Osik JJ, Ritter NJ, Wang S, Shaw JT, Fiser J, Van Hooser SD (2016) Optogenetic spatial and temporal control of cortical circuits on a columnar scale. J Neurophysiol 115:1043–1062.

Rubin DB, Van Hooser SD, Miller KD (2015) The stabilized supralinear network: a unifying circuit motif underlying multi-input integration in sensory cortex. Neuron 85:402–417.

Ruff DA, Alberts JJ, Cohen MR (2016) Relating normalization to neuronal populations across cortical areas. J Neurophysiol 116:1375–1386.

Ruthazer ES, Stryker MP (1996) The role of activity in the development of long-range horizontal connections in area 17 of the ferret. J Neurosci 16:7253–7269.

Sato TK, Hausser M, Carandini M (2014) Distal connectivity causes summation and division across mouse visual cortex. Nat Neurosci 17:30–32.

Sato TK, Haider B, Hausser M, Carandini M (2016) An excitatory basis for divisive normalization in visual cortex. Nat Neurosci 19:568–570.

Sceniak MP, Ringach DL, Hawken MJ, Shapley R (1999) Contrast's effect on spatial summation by macaque V1 neurons. Nat Neurosci 2:733–739.

Self MW, Lorteije JA, Vangeneugden J, van Beest EH, Grigore ME, Levelt CN, Heimel JA, Roelfsema PR (2014) Orientation-tuned surround suppression in mouse visual cortex. J Neurosci 34:9290–9304.

Sengpiel F, Sen A, Blakemore C (1997) Characteristics of surround inhibition in cat area 17. Exp Brain Res 116:216–228.

Simoncelli EP, Heeger DJ (1998) A model of neuronal responses in visual area MT. Vision Res 38:743–761.

Smith GB, Sederberg A, Elyada YM, Van Hooser SD, Kaschube M, Fitzpatrick D (2015) The development of cortical circuits for motion discrimination. Nat Neurosci 18:252–261.

Somers D, Nelson SB, Sur M (1995) An emergent model of orientation selectivity in cat visual cortical simple cells. J Neurosci 15:5448–5465.

Tavazoie SF, Reid RC (2000) Diverse receptive fields in the lateral geniculate nucleus during thalamocortical development. Nat Neurosci 3:608–616.

Tolhurst DJ, Heeger DJ (1997) Comparison of contrast-normalization and threshold models of the responses of simple cells in cat striate cortex. Vis Neurosci 14:293–309.

Turrigiano GG, Nelson SB (2000) Hebb and homeostasis in neuronal plasticity. Curr Opin Neurobiol 10:358–364.

Van den Bergh G, Zhang B, Arckens L, Chino YM (2010) Receptive-field properties of V1 and V2 neurons in mice and macaque monkeys. J Comp Neurol 518:2051–2070.

van Hateren JH, van der Schaaf A (1998) Independent component filters of natural images compared with simple cells in primary visual cortex. Proc Biol Sci 265:359–366.

Van Hooser SD, Heimel JA, Chung S, Nelson SB (2006) Lack of patchy horizontal connectivity in primary visual cortex of a mammal without orientation maps. J Neurosci 26:7680–7692.

Van Hooser SD, Li Y, Christensson M, Smith GB, White LE, Fitzpatrick D (2012) Initial neighborhood biases and the quality of motion stimulation jointly influence the rapid emergence of direction preference in visual cortex. J Neurosci 32:7258–7266.

Vinje WE, Gallant JL (2002) Natural stimulation of the nonclassical receptive field increases information transmission efficiency in V1. J Neurosci 22:2904–2915.

Wang BS, Sarnaik R, Cang J (2010) Critical period plasticity matches binocular orientation preference in the visual cortex. Neuron 65:246–256.

White LE, Fitzpatrick D (2007) Vision and cortical map development. Neuron 56:327–338.

White LE, Coppola DM, Fitzpatrick D (2001) The contribution of sensory experience to the maturation of orientation selectivity in ferret visual cortex. Nature 411:1049–1052.

Williams AL, Singh KD, Smith AT (2003) Surround modulation measured with functional MRI in the human visual cortex. J Neurophysiol 89:525–533.

Zenger-Landolt B, Heeger DJ (2003) Response suppression in v1 agrees with psychophysics of surround masking. J Neurosci 23:6884–6893.

Zoccolan D, Cox DD, DiCarlo JJ (2005) Multiple object response normalization in monkey inferotemporal cortex. J Neurosci 25:8150–8164.

